# HIF-1α regulates mitochondrial function in bone marrow-derived macrophages, but not in tissue-resident alveolar macrophages

**DOI:** 10.1101/2024.10.14.618294

**Authors:** Parker S. Woods, Rengül Cetin-Atalay, Angelo Y. Meliton, Kaitlyn A. Sun, Obada R. Shamaa, Kun Woo D. Shin, Yufeng Tian, Benjamin Haugen, Robert B. Hamanaka, Gökhan M. Mutlu

## Abstract

HIF-1α plays a critical role in shaping macrophage phenotype and effector function. We have previously shown that tissue-resident alveolar macrophages (TR-AMs) have extremely low glycolytic capacity at steady-state, but can shift toward glycolysis under hypoxic conditions. Here, using inducible HIF-1α knockout (*Hif1a^-/-^*) TR-AMs and bone marrow-derived macrophages (BMDMs) and show that TR-AM HIF-1α is required for the glycolytic shift under prolyl hydroxylase inhibition, but is dispensable at steady-state for inflammatory effector function. In contrast, HIF-1α deletion in BMDMs led to diminished glycolytic capacity at steady-state and reduced inflammatory capacity, but higher mitochondrial function. Gene set enrichment analysis revealed enhanced c-Myc transcriptional activity in *Hif1a^-/-^* BMDMs, and upregulation of gene pathways related to ribosomal biogenesis and cellular proliferation. The findings highlight the heterogeneity of HIF-1α function in distinct macrophage populations and provide new insight into how HIF-1α regulates gene expression, inflammation, and metabolism in macrophages.

## Introduction

It is well-established that glycolytic metabolism plays a central role in regulating immune cell effector function.^1–5^ As the field of immunometabolism grows, it has become more apparent that tissue-specific conditions lead to unique metabolic characteristics that often fall outside of the conventional framework ascribed to broader immune cell populations.^6–8^ For instance, while bone marrow-derived macrophages (BMDMs) rely predominantly on glycolysis for proinflammatory processes, we have found glycolysis to be dispensable for inflammation in tissue-resident alveolar macrophages (TR-AMs).^9^ It is likely that the unique environment of the alveolar lumen dictates the metabolic needs of TR-AMs.^10^ Glucose concentrations within the airway are approximately one-tenth of those observed in the blood, and oxygen levels are the highest of any compartment within the human body making oxidative phosphorylation a more efficient means of metabolism for TR-AMs.^6,11^ While TR-AMs do not conduct glycolysis under steady-state conditions, they can be pushed toward a glycolytic phenotype under severe hypoxia such as in cases of acute respiratory distress syndrome (ARDS) where lung oxygenation is impaired.^12^ Further highlighting the role of the alveolar niche in dictating cellular phenotype, BMDMs, peritoneal macrophages, and macrophage precursors transplanted into the airway acquire phenotypical characteristics associated with TR-AM identity.^8,13^ Together, these findings show that TR-AMs occupy a unique environment that negates the need for glycolytic metabolism under steady-state conditions.

It is widely understood that the transcription factor hypoxia-inducible factor 1-alpha (HIF-1α) is a key regulator of glycolysis, particularly at low oxygen concentrations.^14^ Under normoxia, HIF-1α is marked for proteasomal degradation by oxygen-dependent prolyl hydroxylases. In contrast, under hypoxia, prolyl hydroxylase activity is diminished so that HIF-1α accumulates and can enter the nucleus to promote transcriptional adaptation to low oxygen levels. Much of the HIF-1α transcriptional response relates to cell survival under oxygen depleted conditions, including upregulation of genes related to glycolysis, angiogenesis, and cell survival. In macrophages of monocytic origin, HIF-1α functions outside the context of the hypoxic response. HIF-1α has been shown to be a key regulator of macrophage migration and proinflammatory processes.^15–18^ Both HIF-1α and glycolysis are induced in BMDMs treated with lipopolysaccharide (LPS), but whether HIF-1α is required for the observed glycolytic reprograming or just for inflammatory processes remains unknown. Moreover, HIF-1α behaves differently in TR-AMs.^12,19^ As we have shown, hypoxia induces HIF-1α stabilization in TR-AMs, but not in BMDMs.^12^ HIF-1α and its target genes are downregulated in TR-AMs following birth, and this downregulation is required for normal TR-AM maturation and function.^19^ Thus, the full extent for how HIF-1α regulates macrophage function remains unclear.

To gain a more complete understanding for how HIF-1α regulates macrophage metabolism and effector function, we deleted HIF-1α from both BMDMs and TR-AMs and performed metabolic and immunological assays. We found that deletion of HIF-1α in mature TR-AMs had no effect on steady-state metabolic function or gene expression. Phenotypical alterations only became apparent in the presence of prolyl hydroxylase inhibitor FG-4592 where HIF-1α knockout TR-AMs failed to take on a glycolytic phenotype and remained highly susceptible to cell death in the presence of electron transport chain (ETC) inhibitors. Conversely, HIF-1α deletion in BMDMs greatly diminished their glycolytic capabilities at both steady-state and in the presence of FG-4592, sensitizing these cells to ETC inhibitor-induced cell death. HIF-1α knockout BMDMs presented with higher mitochondrial oxygen consumption rates, TCA metabolite levels, and ETC gene and protein expression compared to controls. This was not the case in TR-AMs.

We found that neither HIF-1α nor glycolysis was induced in TR-AMs upon LPS stimulation, and that TR-AMs with HIF-1α knockout exhibited no observable alterations in TCA metabolite levels or cytokine production. In contrast, deletion of HIF-1α in BMDMs led to reductions in LPS-induced glycolysis, but the magnitude of the response did not differ substantially from control BMDMs. HIF-1α knockout BMDMs also exhibited reductions in cytokine production, but presented with higher TCA metabolite levels following LPS treatment. Gene set enrichment analysis of HIF-1α knockout BMDMs revealed the upregulation c-Myc-dependent transcriptional programs, including genes related to ribosomal biogenesis and cellular proliferation. Collectively, these findings highlight the heterogeneity of HIF-1α function in distinct macrophage populations and provide new insight into how HIF-1α regulates inflammation and metabolism in macrophages.

## Results

### HIF-1α deletion alters transcriptome both at baseline and following treatment with a HIF-1α stabilizer in BMDMs but only after HIF-1α stabilizer in TR-AMs

We have recently demonstrated that TR-AMs have a very low glycolytic capacity at steady-state, but can be pushed toward a glycolytic phenotype under hypoxic conditions.^12^ Whether this metabolic reprogramming depends on HIF-1α is unknown. Thus, we generated inducible HIF-1α knockout (*Hif1a^-/-^*) TR-AMs and BMDMs (Figure 1A, D, respectively), and performed RNA-sequencing to determine the role of HIF-1α in transcriptional activity in TR-AMs and BMDMs. Both wildtype control (*Hif1a^+/+^*) and HIF-1α knockout (*Hif1a^-/-^*) macrophages were treated overnight with FG-4592 (a potent prolyl hydroxylase inhibitor, which stabilizes HIF-1α protein) or left untreated. We found that loss of HIF-1α in TR-AMs resulted in minimal changes in gene expression at baseline with only 10 observed significantly differentially expressed genes (DEGs, Log2 fold change ≥ 1 and p≤0.05) (8 upregulated and 2 downregulated) when compared to controls (Figure 1B, Figure S1). Treatment of control (*Hif1a^+/+^*) TR-AMs with FG-4592 led to 953 DEGs (522 upregulated and 431 downregulated). Compared to control TR-AMs, the effect of FG-4592 was greatly diminished in *Hif1a^-/-^* TR-AMs which exhibited 197 DEGs (157 upregulated and 40 downregulated). Heatmap analysis for HIF-1α target genes confirmed that *Hif1a^-/-^* TR-AMs exhibited a greatly diminished response to FG-4592 in a HIF-1α specific manner (Figure 1C).

**Figure 1.**
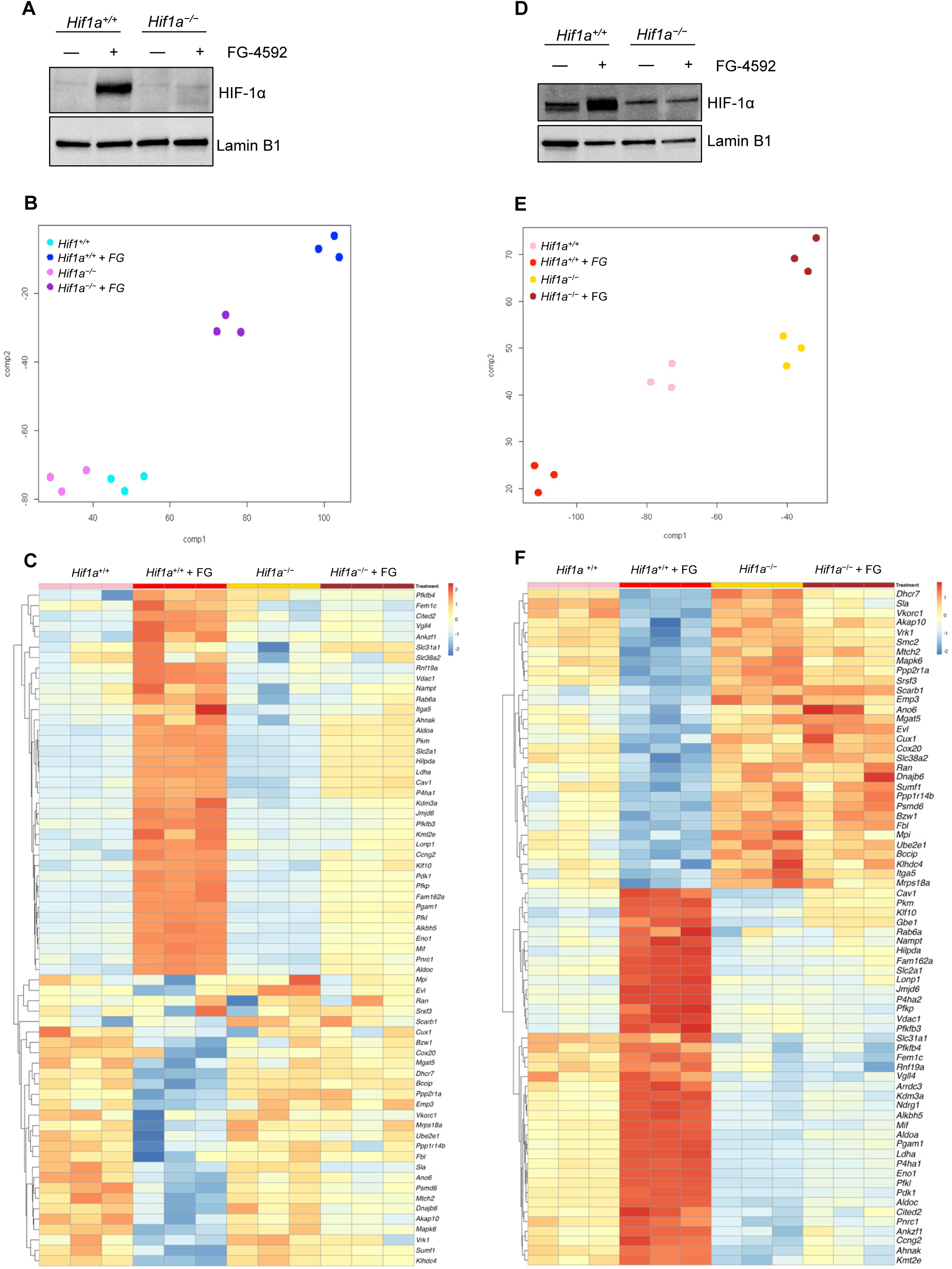
HIF-1α deletion alters transcriptome both at baseline and following treatment with a HIF-1α stabilizer in BMDMs but only after HIF-1α stabilizer in TR-AMs. TR-AMs **(A-C)** and BMDMs Figure 1. HIF-1α deletion alters tr were cultured overnight (16h) in the presence or absence of FG-4592 (25µM). (A, D) Western blot analysis of nuclear extracts to verify successful HIF-1α knockout in our transgenic system. (B, E) Clustering of samples based on RNA-seq gene expression profiles using t-SNE projection (C, F) Heatmap analysis of CHEA (ChIP-X Enrichment Analysis) HIF-1α transcription factor targets.

In contrast to TR-AMs, HIF-1α deletion resulted in significant alterations at baseline in BMDMs with 305 DEGs (121 upregulated and 184 downregulated). After FG-4592 treatment, only 126 DEGs (60 upregulated and 66 downregulated) were observed in *Hif1a^-/-^* BMDMs compared to 694 DEGs (323 upregulated and 371 downregulated) in control BMDMs (Figure S1B). Heatmap analysis for HIF-1α target genes further demonstrated that *Hif1a^-/-^* BMDMs exhibit significant basal alterations in gene expression and a substantial reduction in their response to FG-4592 in a HIF1α-specific manner (Figure 1F).

Collectively, these data validate our previous findings documenting that HIF-1α is dispensable in TR-AMs under steady-state conditions. These data also support the current literature pointing to HIF-1α as a key regulator of normal BMDM transcriptional response and function.

### HIF-1α deletion broadly impairs glycolysis in BMDMs but only impairs TR-AM glycolysis following HIF-1α stabilizer treatment

We next sought to assess the functional consequences of HIF-1α deletion in our two macrophage populations by evaluating overall glycolytic fitness. To do so, we performed a glycolysis stress test and measured glycolysis, assessed by extracellular acidification rate (ECAR), after overnight (16 hours) treatment with FG-4592. Loss of HIF-1α did not affect baseline glycolytic rate in TR-AMs, as *Hif1a^-/-^* TR-AMs did not exhibit any changes in their glycolytic rate compared to control TR-AMs, which already had low glycolysis (Figure 2A and B). As expected, FG-4592 induced an increase in glycolytic rate in control *Hif1a^+/+^* TR-AMs (Figure 2A and B). However, after treatment with FG-4592, *Hif1a^-/-^* TR-AMs had significantly lower glycolysis compared to treated controls (Figure 2A, B). Correlating with the low glycolytic rate, both *Hif1a^+/+^* and *Hif1a^-/-^* TR-AMs had lower expression of glycolytic genes and proteins (Figure 2C, D). Consistent with the role of HIF-1α in glycolytic gene expression, *Hif1a^-/-^* TR-AMs treated with FG-4592 had reduced induction of glycolytic gene and protein expression (Figure 2C, D). We have previously demonstrated TR-AMs are exquisitely sensitive to inhibition of the ETC and that hypoxic or pharmacological induction of HIF-1α could rescue TR-AMs treated with mitochondrial ETC inhibitors from cell death.^12^ In support of these findings, FG-4592 became ineffective in rescuing *Hif1a^-/-^* TR-AMs from ETC inhibitor-induced cell death (Figure 2E).

**Figure 2.**
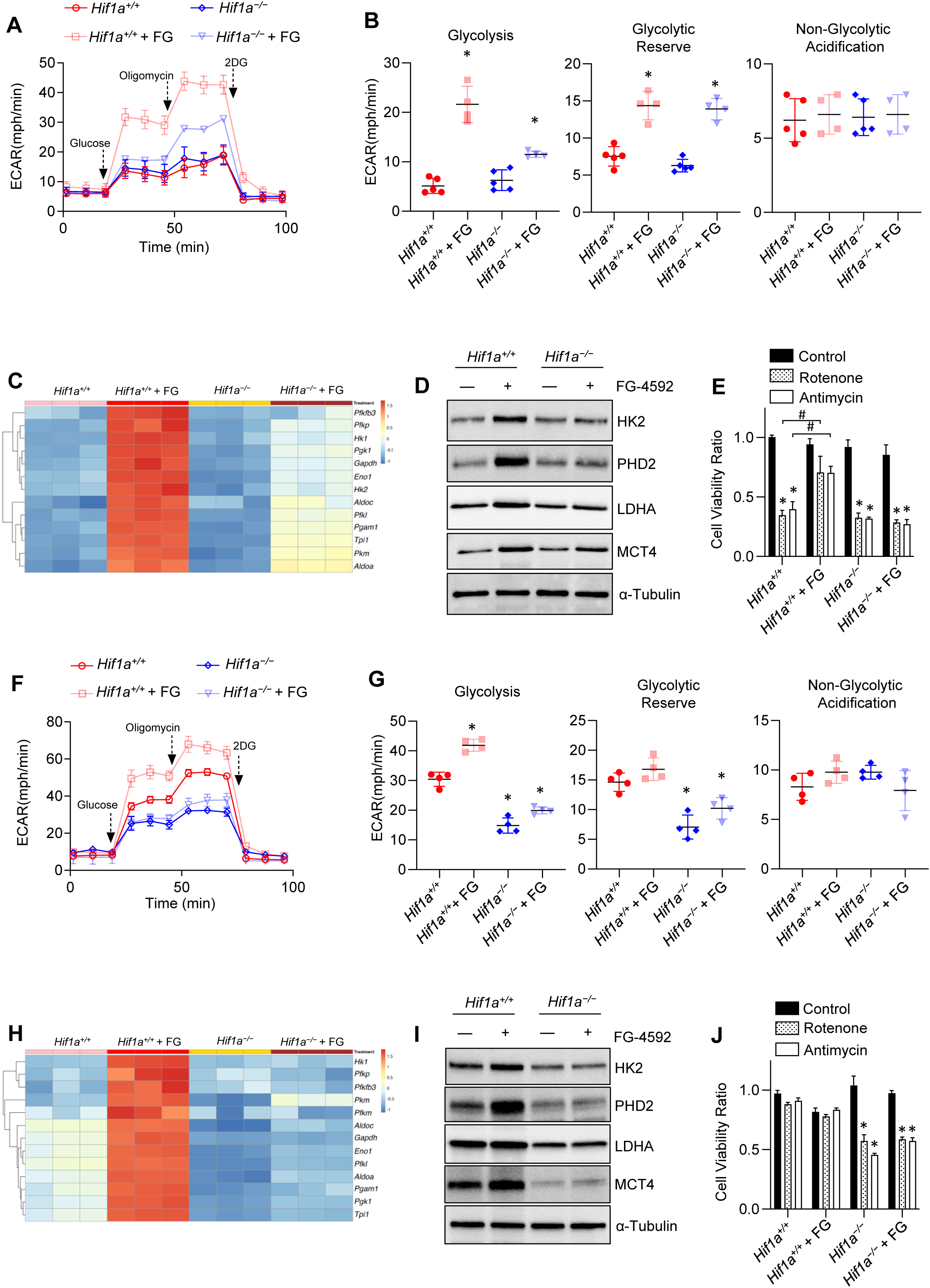
HIF-1α deletion broadly impairs glycolysis in BMDMs but only impairs TR-AM glycolysis following HIF-1α stabilizer treatment. TR-AMs **(A-E)** and BMDMs **(F-J)** were cultured overnight (16h) in the presence or absence of FG-4592 (25µM). **(A, F)** Using Seahorse XF24 analyzer, glycolysis was measured as extracellular acidification rate (ECAR). Macrophages were treated in succession with glucose, oligomycin (ATP synthase inhibitor) and 2-deoxyglucose (2-DG) (inhibitor of hexokinase 2, or glycolysis). **(B, G)** Scatter plots quantifying glycolytic parameters. Data represent at least 3 independent experiments (n=4 separate wells per group). All groups compared against *Hif1a^+/+^* no treatment group with significance determined by two-way ANOVA with Bonferroni’s post test. All error bars denote mean ± SD. *, p < 0.05. **(C, H)** Heatmap of glycolytic specific gene expression in macrophages. **(D,I)** Western blot analysis of macrophage lysates to assess glycolytic protein expression. **(E, J)** Macrophages were first treated with FG-4592 for 8h then treated overnight with mitochondrial inhibitors (500nM Antimycin or 500nM Rotenone). Sulforhodamine B assay was performed to measure cytotoxicity. Graphs represent cell viability compared to control, *Hif1a^+/+^* no treatment group. Data represent at least 3 independent experiments (n=3 per group). All groups compared against *Hif1a^+/+^* no treatment group with significance determined by two-way ANOVA with Bonferroni’s post test. All error bars denote mean ± SD. *, p < 0.05 signifies reduced cell viability compared to *Hif1a^+/+^* no treatment group; #, p < 0.05 signifies enhanced viability compared to *Hif1a^+/+^* Rotenone and Antimycin treated groups.

Our previous work demonstrated that BMDMs have elevated basal levels of HIF-1α compared with TR-AMs and that hypoxia does not increase the glycolytic rate of these cells. Here, we observed that control BMDMs treated with FG-4592 presented with higher ECAR compared to untreated controls (Figure 2F). However, this increase in BMDMs was more modest (∼30%) compared to the 4-fold increase in ECAR following FG-4592 in TR-AMs (Figure 2A). Glycolytic gene and protein expression were also higher in control BMDMs treated with FG-4592 (Figure 2H, I). Glycolytic rates were significantly lower in *Hif1a^-/-^* BMDMs, and no observable alterations in the ECAR response were seen with FG-4592 treatment. Glycolytic gene and protein expression were also lower in *Hif1a^-/-^* BMDMs compared to controls and was unaffected by FG-4592 treatment (Figure 2H, I). BMDMs are resistant to ETC inhibition-induced death at baseline but they became sensitized after the loss of HIF-1α (Figure 2J). These data demonstrate that deletion of HIF-1α results in impaired glycolysis in both TR-AMs and BMDMs. Consistent with our previous findings that TR-AMs contain very little expression of HIF-1α protein at baseline, we found that loss of HIF-1α does not affect glycolysis in TR-AMs at baseline and affects glycolysis only after treatment with FG-4592. BMDMs, which have high basal levels of HIF-1α exhibited reduction in glycolysis following the loss of *Hif1a* both at baseline and after FG-4592 suggesting that HIF-1α regulates metabolic function in BMDMs both in normoxia as well as during prolyl hydroxylase inhibition.

### HIF-1α deletion boosts mitochondrial function in BMDMs, but has limited impact on TR-AM mitochondrial function

The effects of HIF-1α deletion on glycolytic function and inflammation have been well-described in BMDMs, but little is known about how HIF-1α deletion impacts mitochondrial function in macrophages.^15^ To determine how HIF-1α regulates mitochondrial function, we performed a mitochondrial stress test and found that *Hif1a^-/-^* TR-AMs exhibited basal oxygen consumption rates (OCRs) comparable to untreated controls (Figure 3A, B). FG-4592 treated control *Hif1a^+/+^* TR-AMs exhibited reduced basal OCR, suggesting a shift toward glycolytic ATP production, but they were still able to reach maximal oxygen consumption under FCCP stimulation. This glycolytic shift was confirmed by the reciprocal change in ECAR measurement where control *Hif1a^+/+^* TR-AMs treated with FG-4592 no longer exhibited reduction in ECAR following rotenone and antimycin A injection whereas *Hif1a^-/-^* TR-AMs treated with FG-4592 showed a reduction in ECAR (Figure S2A). Carbonic acid derived from mitochondrial CO_2_ can be measured as the ECAR during mitochondrial stress test.^20^ The reduction in *Hif1a^-/-^* TR-AMs ECAR after administration of rotenone and antimycin A support the notion that the ECAR is mitochondria-derived carbonic acid, while a lack of ECAR responsiveness in control *Hif1a^+/+^* TR-AMs signifies glycolytically produced acid. Interestingly, *Hif1a^-/-^* TR-AMs did present with higher maximal respiration/spare capacity compared to controls, although GC-MS analysis did not reveal any alterations in *Hif1a^-/-^* TR-AM TCA metabolites compared to controls (data not shown). This suggests that the changes in *Hif1a^-/-^* TR-AMs’ maximal oxygen consumption have little functional consequence under normal, non-stressed conditions.

**Figure 3.**
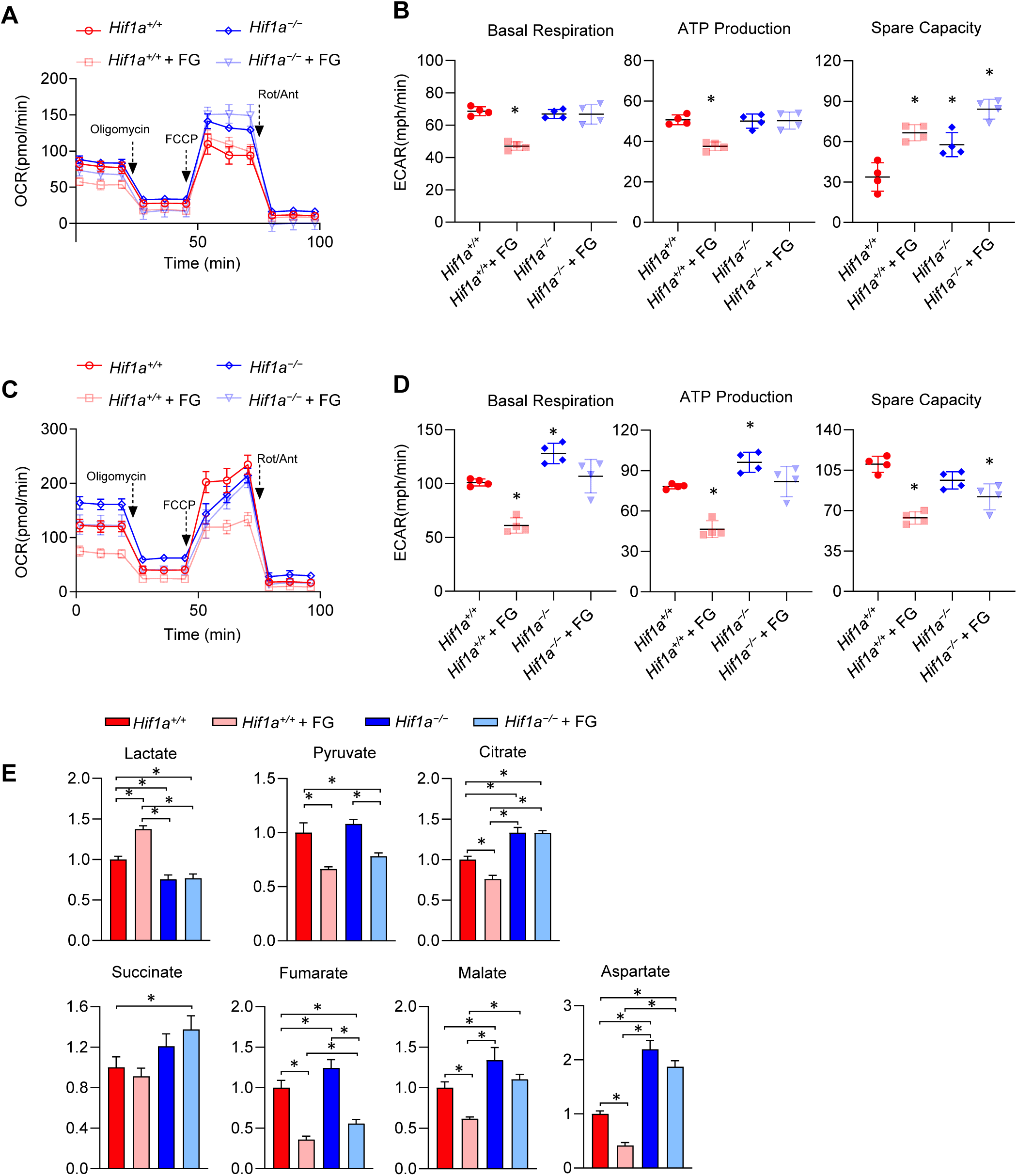
HIF-1α deletion boosts mitochondrial function in BMDMs, but has limited impact on TR-AM mitochondrial function. TR-AMs **(A, B)** and BMDMs **(C-E)** were cultured overnight (16h) in the presence or absence of FG-4592 (25µM). **(A, C)** Mitochondrial stress test to measure oxygen consumption rate (OCR) using Seahorse XF24 in TR-AMs and BMDMs. Macrophages were treated sequentially with oligomycin (ATP synthase inhibitor), FCCP (uncoupler) and Rotenone/Antimycin A (complex I and III inhibitor, respectively). **(B, D)** Interleaved scatter plots quantifying mitochondrial respiration parameters. Data represents at least 3 experiments (n=4 separate wells per group). Mitochondrial parameters were compared against *Hif1a^+/+^* no treatment group and significance was determined by two-way ANOVA with Bonferroni’s post test. **(E)** GC-MS metabolite analysis of BMDMs. All groups compared against *Hif1a^+/+^* no treatment group with significance determined by two-way ANOVA with Bonferroni’s post test. All error bars denote mean ± SD. *, p < 0.05.

Unlike TR-AMs, control BMDMs treated with FG-4592 showed diminished OCR across all parameters compared to untreated controls. HIF-1α deletion in BMDMs led to significantly higher basal respiration and ATP production compared to controls and the effect of FG-4592 on OCR was greatly diminished (Figure 3C, D). In agreement with the OCR data, GC-MS analysis showed that key TCA cycle metabolites were significantly elevated in *Hif1a^-/-^* BMDMs, and that the levels of these metabolites were less responsive to FG-4592 treatment compared to controls (Figure 3E).

To bolster our mitochondrial stress test and GC-MS data, we performed western blot analysis using an antibody cocktail that recognizes components of each of the respiratory complexes. *Hif1a^-/-^* TR-AMs did not exhibit any significant alterations in ETC complex protein expression compared to controls, and FG-4592 had no effect on ETC protein expression across all groups (Figure 4A). Heatmap analysis of respiratory subunit gene expression revealed that FG-4592 reduced the expression of OXPHOS-related genes in control *Hif1a^+/+^* TR-AMs, and that this response was not observed in *Hif1a^-/-^* TR-AMs treated with FG-4592 (Figure 4B). Unlike TR-AMs, HIF-1α deletion in BMDMs led to significant increases in protein expression for ETC complexes I-IV (Figure 4C). FG-4592 did not alter ETC protein expression in control or *Hif1a^-/-^* BMDMs. Like control TR-AMs, FG-4592 reduced OXPHOS-related gene expression in control BMDMs, but did not affect OXPHOS-related gene expression in *Hif1a^-/-^* BMDMs (Figure 4D). *Hif1a^-/-^* BMDMs also had higher basal OXPHOS-related gene expression than controls which correlates with their protein expression data.

**Figure 4.**
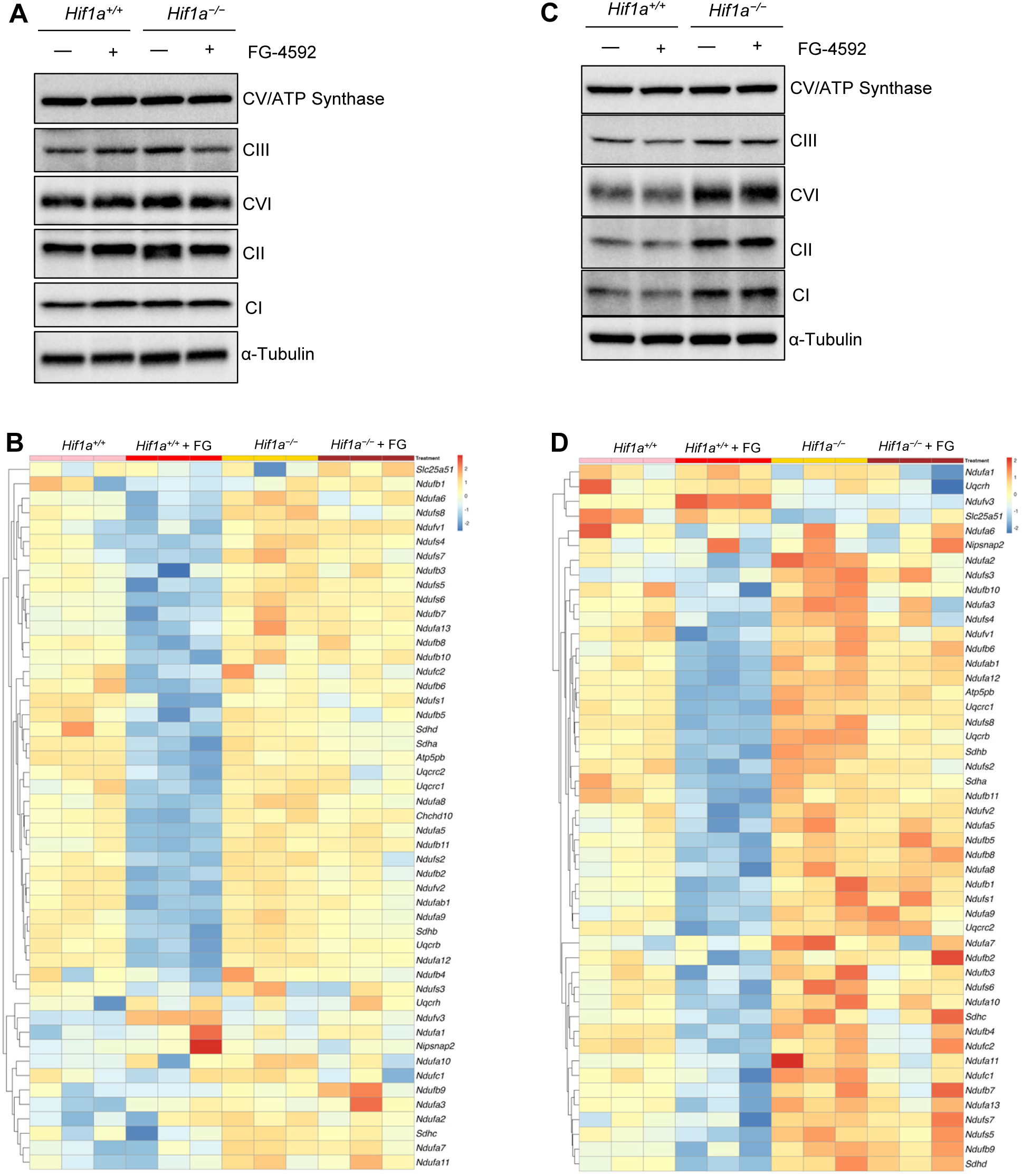
HIF-1α deletion results in increased Mitochondrial gene and protein expression in BMDMs, but not TR-AMs. TR-AMs **(A-B)** and BMDMs **(C-D)** were cultured overnight (16h) in the presence or absence of FG-4592 (25µM). **(A, C)** Western blot analysis of macrophage lysates to assess electron transport chain protein expression. **(B, D)** Heatmap of gene expression related to oxidative phosphorylation.

Taken together, these data suggest that HIF-1α serves as a negative regulator of mitochondrial function in BMDMs under steady-state conditions. Moreover, HIF-1α stabilization reduces OXPHOS-related gene expression in both TR-AMs and BMDMs, but this does not correlate with the reduction in OXPHOS protein expression, which remains unchanged in the presence of FG-4592.

### HIF-1α deletion impairs inflammatory capacity in BMDMs, but not in TR-AMs

The central role of HIF-1α in macrophage inflammation has been well-documented.^21^ Here, we wanted to determine more thoroughly the metabolic and inflammatory alterations that occur in the absence of HIF-1α. We and others have shown the immediate upregulation of glycolysis upon LPS exposure in BMDMs.^9,22,23^ To determine if the increase in glycolytic output was mediated by HIF-1α, we treated *Hif1a^-/-^* BMDMs and controls with LPS to observe changes in ECAR over a five-hour period. Figure 5A shows that *Hif1a^-/-^* BMDMs still respond to LPS by immediately upregulating glycolysis. This response is attenuated compared to controls, which is likely a result of *Hif1a^-/-^* BMDMs having lower glycolytic gene and protein expression (Figure 2H, I). While basal ECAR was lower in *Hif1a^-/-^* BMDMs compared to controls, there was no change in the magnitude of glycolytic induction when the ECAR was normalized to percent change (ECAR %) from baseline (Figure 5B). This would suggest that HIF-1α is not required for the enhanced glycolysis observed under LPS stimulation. In agreement with these data, siRNA knockdown of HIF-1α in BMDMs did not alter glycolytic responsiveness to LPS (Figure S3A, B). The glycolytic rates of *Hif1a^-/-^* and control TR-AMs remained unresponsive to LPS injection, which is in line with our previous findings (Figure S3C) ^9^.

**Figure 5.**
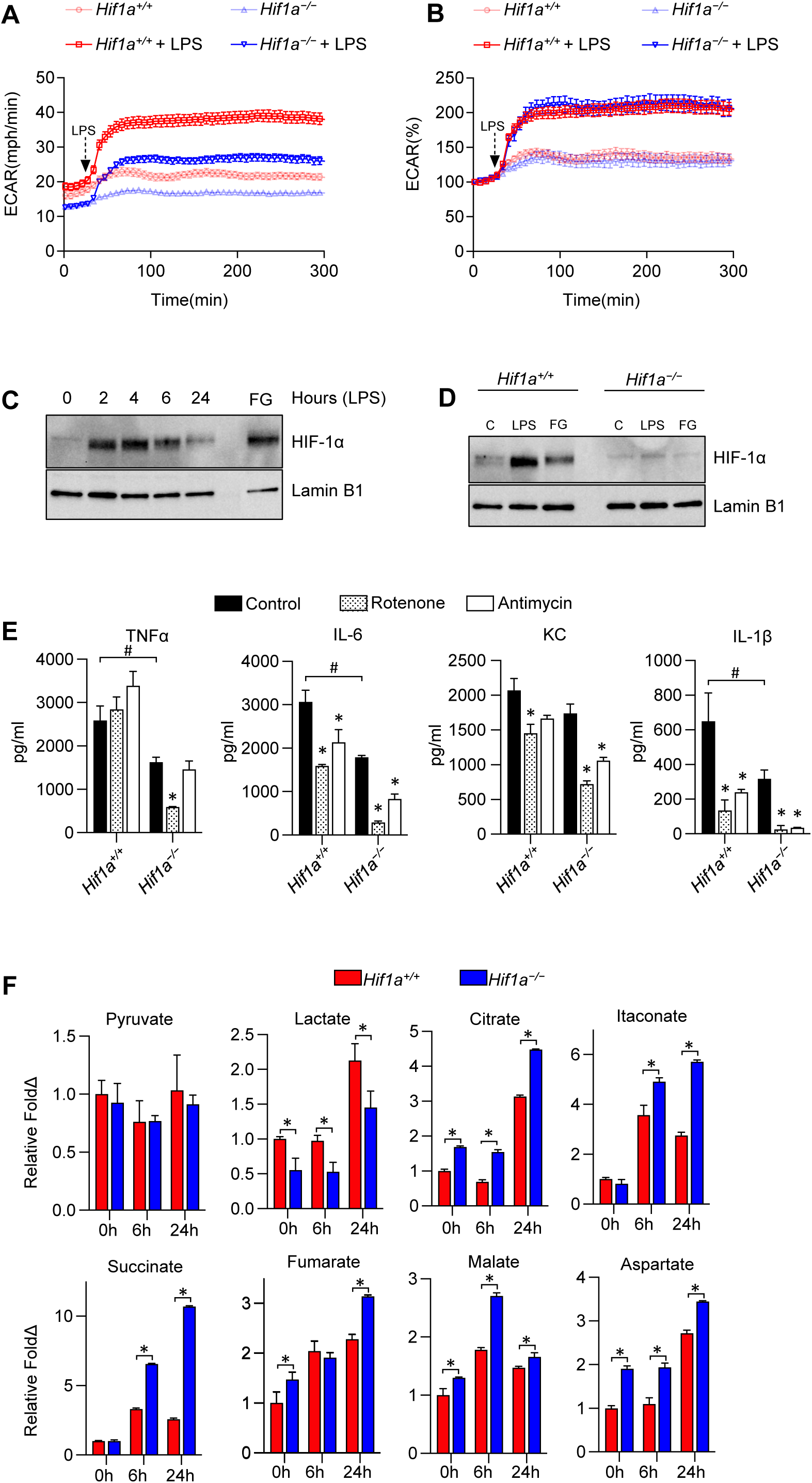
HIF-1α deletion impairs inflammatory capacity in BMDMs, but not TR-AMs. **(A,B)** ECAR was measured following acute LPS injection (final concentration: 20 ng/ml) in BMDMs. ECAR data represented both as **(A)** raw values and **(B)** % change from baseline. Data represent at least 3 independent experiments (n=4 separate wells per group). Error bars denote mean ± SEM to allow for easier visualization of line graphs. **(C)** Western blot analysis assessing nuclear localization of HIF-1α over 24 hour LPS (20ng/ml) time course. DMOG was used as positive HIF-1α control. **(D)** Western blot at 4 hours verifying our protein of interest is in fact HIF-1α. **(E)** BMDMs were treated with LPS (20ng/ml) in the presence or absence of Antimycin A (20nM) or Rotenone (20nM) for 6 hours and secreted cytokines were measured via ELISA. Significance determined by two-way ANOVA with Bonferroni’s post test. All error bars denote mean ± SD. Significance markings represented in relation to *Hif1a^+/+^* control group with #, p < 0.05. *, p < 0.05 denotes significance in relation to control groups within respective genotypes. **(F)** GC-MS metabolite analysis of BMDM LPS (20ng/ml) time course. Significance determined by two-way ANOVA with Bonferroni’s post test. All error bars denote mean ± SD. Significance markings represented in relation to *Hif1a^+/+^ vs Hif1a^−/−^* at a given timepoint with *, p < 0.05.

We next examined the nuclear localization of HIF-1*⍺* in macrophages following LPS treatment. Several labs have shown HIF-1α protein stabilization in the presence of LPS, but the exact time-course of stabilization remains unclear. We found that nuclear stabilization of HIF-1α in BMDMs occurs early in exposure, peaks around 4 hours, and returns to near baseline levels by 24 hours (Figure 5C). We confirmed the loss of HIF-1α protein in our *Hif1a^-/-^* BMDMs under LPS stimulation by verifying the lack of nuclear expression using western blot (Figure 5D) and immunofluorescence (Figure S4). Moreover, we have only observed HIF-1α induction in TR-AMs under hypoxia or pseudohypoxia (prolyl hydroxylase inhibition with FG-4592), but we have not seen HIF-1α induction/stabilization in TR-AMs under inflammatory stimuli. Immunofluorescence and western blot confirmed these observations where HIF-1α was only observed in the presence of FG-4592 in TR-AMs, but not in the presence of LPS (Figure S5, S6). Poly (I:C), a potent TLR3 antagonist, also failed to induce HIF-1α in TR-AMs.

We next wanted to determine the effect of HIF-1α deletion on macrophage cytokine production and inflammatory metabolite levels. Compared to control BMDMs, *Hif1a^-/-^* BMDMs secreted less TNFα, IL-6, and IL-1β in response to LPS and also exhibited reduced cytokine secretion in the presence of an ETC inhibitor (Figure 5E). By contrast, TR-AM cytokine secretion was unperturbed by HIF-1α deletion, and both *Hif1a^-/-^* and control TR-AMs were equally susceptible to ETC inhibitor-induced cytokine impairments. (Figure S7A). Next, BMDMs and TR-AMs were treated with LPS for 6 and 24 hours and the cellular extracts were analyzed via GC-MS. As expected, *Hif1a^-/-^* BMDMs had reduced lactate production in response to LPS (Figure 5F). While pyruvate levels remained unchanged, all other TCA cycle metabolites were significantly elevated in *Hif1a^-/-^* BMDMs treated with LPS compared to controls. Most notably, succinate, citrate, and itaconate, which are known regulators of macrophage inflammation, were robustly elevated in *Hif1a^-/-^* BMDMs. HIF1α deletion in TR-AMs had negligible effects on LPS-induced metabolite levels (Figure S7B). Malate was the only TCA metabolite that was elevated in *Hif1a^-/-^* TR-AMs compared to controls. These data indicate that HIF-1α deletion in BMDMs leads to reduced cytokine production and elevated TCA metabolite levels in response to LPS.

### HIF-1α deletion in BMDMs alters c-Myc transcriptional activity

Thus far, we have demonstrated that HIF-1α deletion is required for TR-AM glycolytic adaptation under prolyl hydroxylase inhibition, but otherwise HIF-1α is dispensable for TR-AM inflammatory processes under steady-state conditions. This is not the case in BMDMs where HIF-1α deletion broadly impacts metabolism and effector function. More specifically, *Hif1a^-/-^* BMDMs have reduced glycolytic and inflammatory capabilities but have enhanced mitochondrial function. By performing GO-BP enrichment analysis, we sought to identify regulatory pathways within our RNA-seq dataset that could explain the altered phenotype in *Hif1a^-/-^* BMDMs. Predictably, GO-BP analysis revealed suppression of biological processes related to hypoxia and oxygen responsiveness in *Hif1a^-/-^* BMDMs (Figure 6A). HIF-1α deletion also led to the activation of biological processes related to ribosome biogenesis, ribosomal RNA processing, and cell growth, all of which can linked to enhanced c-Myc activities.^24^ It is well-known that c-Myc promotes cell proliferation and often works diametrically in relation to HIF-1α.^25^ Thus, we queried c-Myc target genes to assess c-Myc transcriptional activity and found that *Hif1a^-/-^* BMDMs had higher gene expression for the majority of c-Myc targets (Figure 6B), with several of the genes being related to enhanced translational activity and cell proliferation (*Cdkn1a* and *Pcna*). Moreover, the only c-Myc targets that were downregulated in *Hif1a^-/-^* compared to controls were genes related to glycolysis, and these genes have been identified as shared c-Myc and HIF-1α targets (Figure 6C). This observation supports our data that HIF-1α, and not c-Myc, is the primary regulator of glycolysis in BMDMs.

**Figure 6.**
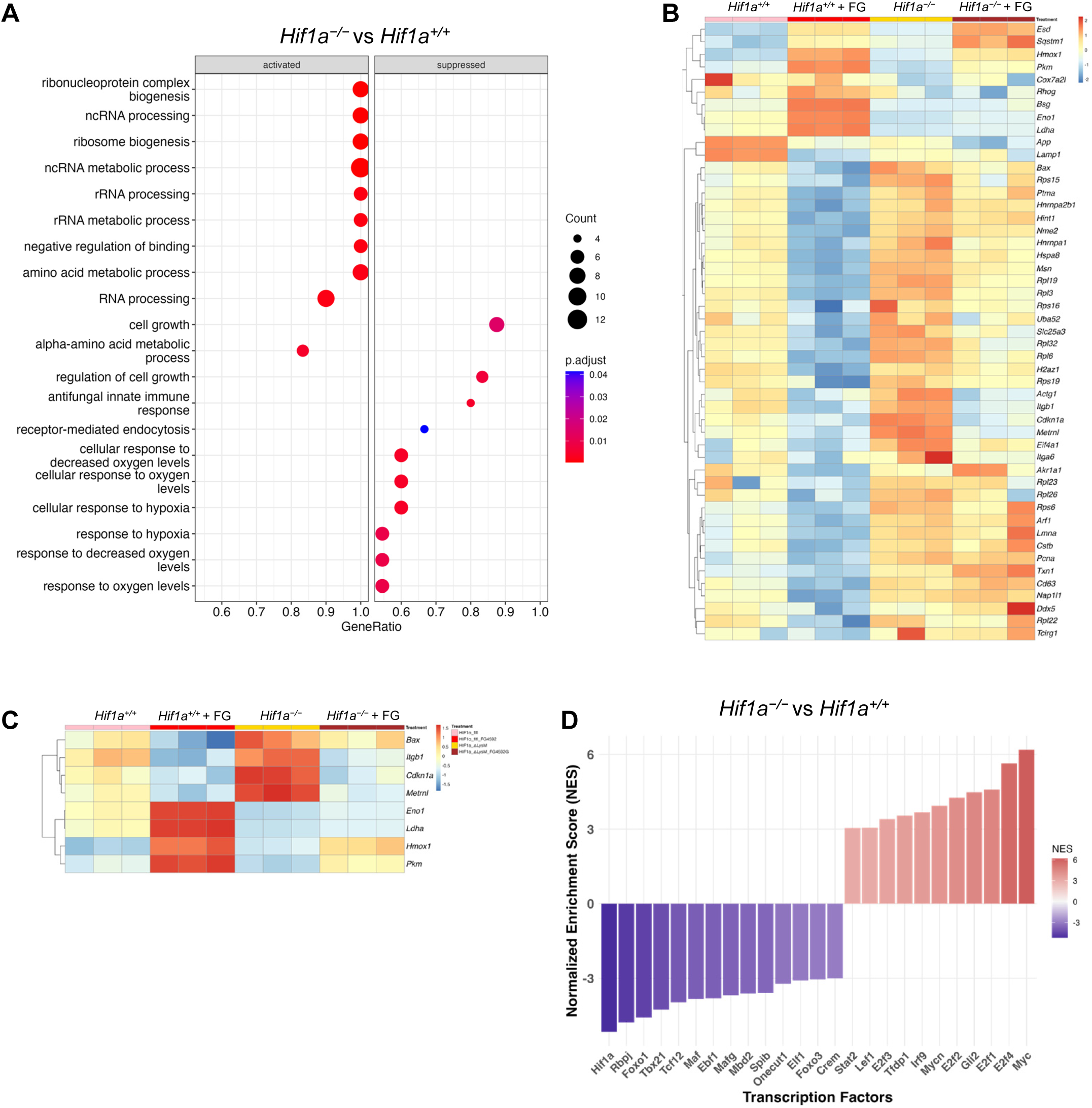
HIF-1α deletion in BMDMs alters c-Myc transcriptional activity. **(A)** GO-BP enrichment analysis of gene sets comparing *Hif1a^−/−^* vs *Hif1a^+/+^* BMDMs at baseline. **(B)** Heatmap of significantly expressed c-Myc target genes and **(C)** Heatmap of shared target genes of c-Myc and HIF-1α comparing *Hif1a^+/+^* and *Hif1a^−/−^* BMDMs ± FG-4592 (25µM) for 16h. All represented DEGs had gene expression abundances exceeding 200 counts. **(D)** Transcription factor enrichment was computed using the DoRothEA R tool with significant differentially expressed genes (logFC ≤ 1, p ≤ 0.05) for *Hif1a^−/−^* vs *Hif1a^+/+^* BMDMs comparisons at baseline. The color legend indicates TF activity.

Given the upregulation of c-Myc target genes in our *Hif1a^-/-^* BMDMs, we next took a non-biased approach using the DoRothEA mouse regulon transcription factor (TF) enrichment computation to estimate the activity of TFs in gene expression data, to understand how TFs drive gene expression changes in a given biological context.^26^ Not surprisingly, the HIF-1α regulon genes had the most negative enrichment score in our *Hif1a^-/-^* BMDMs. In contrast, Myc was the most positively enriched regulatory interaction with its targets in *Hif1a^-/-^* BMDMs. These data support the notion that HIF-1α is a negative regulator of c-Myc transcriptional activity. Thus, with the deletion of HIF-1α in BMDMs, it is possible that c-Myc shifts BMDMs toward a mitochondrial dominated metabolism that favors proliferation/cellular maintenance rather than a glycolytic phenotype that is poised for robust inflammatory effector function.

## Discussion

Macrophages are highly plastic in their responses to immune stimuli. Their proinflammatory processes are necessary for pathogen clearance, while their pro-resolving properties are required in tissue repair and disease resolution. We now know that tissue-specificity is a key determinant for macrophage function. The respiratory tract is the primary portal for pathogen entry into the human body, and acute respiratory distress syndrome (ARDS) caused by influenza virus and SARS-CoV-2 causes significant mortality annually. TR-AMs play a critical role in mediating the response to these viruses as loss of TR-AMs during ARDS is associated with more severe disease and higher morbidity and mortality.^27–29^ Moreover, recruitment of non-resident macrophages to the lung during ARDS is thought to worsen outcomes.^30,31^ Thus, understanding fundamental macrophage processes, such as metabolic adaptation to immune stimuli and local tissue conditions could ultimately lead to new therapeutic strategies to attenuate disease.

We have previously shown that TR-AMs have very low glycolytic activity under steady-state conditions and that glycolysis is dispensable for LPS-induced inflammation in these cells.^9^ We then went on to show that TR-AMs could undergo glycolytic reprogramming at low oxygen concentrations, and that HIF-1α was likely driving this metabolic switch. Treating influenza-infected mice with prolyl hydroxylase inhibitor, FG-4592, promoted TR-AM survival and reduced lung injury.^12^ Together, these previous findings demonstrate that HIF-1α induction in TR-AMs provides a mechanism for cell survival in an injured, hypoxic lung. In this study, we wanted to definitively identify HIF-1α as the driver of glycolytic adaptation in TR-AMs and to rule out HIF-1α as a mediator of TR-AM inflammation. Moreover, we wanted to provide new insight into the central role of HIF-1α in BMDM effector function. To do so, we generated HIF-1α knockout macrophages and found that HIF-1α is not required for steady-state TR-AM function, but it is only required for glycolytic adaptation under prolyl hydroxylase inhibition. RNA-seq analysis revealed *Hif1a^-/-^* TR-AMs had only 10 DEGs compared to controls at steady-state, but 610 DEGs were observed between control and *Hif1a^-/-^* TR-AMs treated with FG-4592. These data were in line with glycolysis stress test and glycolytic protein expression data, and ETC inhibitor-induced cytotoxicity experiments in which *Hif1a^-/-^* TR-AMs failed to undergo glycolytic adaptation in the presence of FG-4592, and thus remained highly susceptible to ETC inhibitor-induced cell death. These data suggest that HIF-1α plays only a minor role in steady-state, terminally differentiated TR-AMs, which is in agreement with the findings that HIF-1α target genes are significantly downregulated in TR-AMs during postnatal maturation.^19^ In contrast, *Hif1a^-/-^* BMDMs exhibited significant alterations at steady-state, presenting with 305 DEGs compared to controls, and the differences increased to 1022 DEGs when treated with FG-4592 (*Hif1a^+l+^* + FG-4592 vs *Hif1a^-/-^* + FG-4592). These data are in line with previous work describing HIF-1α regulation of basal glycolytic metabolism in mouse peritoneal macrophages.^15^ *Hif1a^-/-^* BMDMs also exhibited decreased glycolytic output and protein expression and became susceptible to ETC inhibition-induced cell death. FG-4592 treatment had no functional effect on glycolytic parameters in *Hif1a^-/-^* BMDMs. Overall, our findings demonstrate that *Hif1a^-/-^* BMDMs more closely resemble wildtype/control TR-AMs with regards to glycolytic phenotype, and that HIF-1α is required for normal BMDM metabolic function.

When assessing mitochondrial function in our *Hif1a^-/-^* macrophages, we found that TR-AMs had minimal alterations compared to controls. *Hif1a^-/-^* TR-AMs only showed a slight elevation in spare mitochondrial capacity, but no change in basal respiration, ATP production, or TCA cycle metabolite levels. In contrast, *Hif1a^-/-^* BMDMs maintained higher basal oxygen consumption rates and ATP production, and OCR was less affected by FG-4592 treatment under mitochondrial stress test assessment. Interestingly, reciprocal ECAR measurements, while generally lower in *Hif1a^-/-^* BMDMs, remained fairly unresponsive to mitochondrial inhibitors (rotenone and antimycin A). This suggests that *Hif1a^-/-^* BMDMs still maintain the ability to produce significant amounts of glycolysis-derived lactate under ETC inhibition, but this does not appear to be sufficient to overcome ETC inhibitor-induced cellular cytotoxicity. It may be that in the absence of HIF-1α other transcription factors aid in the basal maintenance of BMDM glycolysis, but that HIF-1α is required for a fully functional glycolytic phenotype.^32–35^ This is not the case in untreated TR-AMs where their reciprocal ECAR measurements return to basal levels regardless of HIF-1α deletion suggesting that acidification measurements represent mitochondrial-derived CO_2_.^20^ Only when wildtype *Hif1a^+l+^* TR-AMs are treated with FG-4592 does their reciprocal ECAR measurements become unresponsive to rotenone and antimycin A suggesting a dominant glycolytic phenotype. HIF-1α deletion in BMDMs also resulted in higher TCA metabolite levels that were less responsive to FG-4592 treatment. These findings correlated with ETC protein and gene levels where *Hif1a^-/-^* BMDMs had significantly higher ETC expression. HIF-1α deletion had no impact on ETC expression at the protein level in TR-AMs. FG-4592 treatment led to downregulation of OXPHOS genes in wildtype TR-AMs and BMDMS, but this effect was lost when HIF-1α was deleted. This suggests that HIF-1α is responsible for the transcriptional downregulation of OXPHOS genes under prolyl hydroxylase inhibition regardless of macrophage origin. Based on these data, it is possible that loss of HIF-1α in BMDMs enhances mitochondrial capabilities as compensatory mechanism for deficiencies in glycolysis or that the presence of HIF-1α serves as negative regulator of mitochondrial function. As a result, *Hif1a^-/-^* BMDM metabolism resembles that of wildtype TR-AMs in their glycolytic deficiency and enhanced mitochondrial metabolism.

It is well-documented that HIF-1α is a key mediator of inflammation in macrophages of monocytic origin and that glycolytic flux increases immediately following proinflammatory immune stimulus exposure.^9,15,22,23^ Here we show that HIF-1α is not responsible for the increased glycolytic flux under LPS stimulation in BMDMs. *Hif1a^-/-^* BMDMs had lower basal ECAR rates under LPS stimulus, but the magnitude of the response was comparable to wildtype BMDMs. Moreover, siRNA knockdown in wildtype BMDMs had no impact on basal ECAR or the glycolytic ECAR flux in response to LPS. This suggests that the increase of glycolytic flux in response to LPS is independent of HIF-1α. Although LPS induces glycolytic gene expression hours after treatment, it is likely that these alterations have little impact on the immediate alterations in glycolytic flux. It is thus more likely that LPS increases the activity of rate-limiting enzymes in the glycolytic pathway, such as PFKFB3 or PKM2, to enhance glycolytic flux.^16,36,37^ We also observed broad decreases in cytokine production in *Hif1a^-/-^* BMDMs in response to LPS. Others have shown that HIF-1α deletion results in deficiencies related to IL-1β production, and that HIF-1α activates NF-κB.^16,18,38,39^ Our findings indicate that broad reductions in cytokine production are likely a result of glycolytic deficiency in *Hif1a^-/-^* BMDMs. Alternatively, in the absence of HIF-1α, BMDMs have elevated TCA cycle metabolite production in response to LPS, including the immunoregulatory metabolite itaconate, which may be dampening the immune response.^40–43^ Similarly, our data indicate that *Hif1a^-/-^* BMDMs have elevated citrate levels both at steady-state and following LPS treatment. Aside from its traditional role in the TCA cycle, citrate can be exported from the mitochondria to generate acetyl-CoA, which can be used for de novo fatty acid synthesis or epigenetic modifications.^44–46^ Others have reported that anti-inflammatory macrophage function relies predominantly on mitochondrial metabolic pathways, which would be in agreement with the enhanced mitochondrial function and reduced proinflammatory cytokine production observed in our *Hif1a^-/-^* BMDMs.^47^

We observed no changes in TR-AM inflammatory capabilities upon HIF-1α deletion. ETC inhibition greatly reduces cytokine production in TR-AMs, but these reductions can be overcome through pretreatment with hypoxia^12^. These findings are aligned with our data showing that unlike BMDMS, LPS treatment does not result in HIF-1α translocation to the nucleus in TR-AMs. Zhu and colleagues have shown that Poly (I:C) or influenza infection upregulate HIF-1α in TR-AMs, but we were not able to replicate these results.^48,49^ We find that only hypoxia or pharmacological inhibition of prolyl hydroxylases induces HIF-1α in TR-AMs. One explanation for the observed differences is that Zhu and colleagues cultured TR-AMs in the presence of GM-CSF. GM-CSF is important to TR-AM function *in vivo*, but it induces proliferation of TR-AMs in culture.^50,51^ Within the alveolus, epithelial-derived soluble mediators and direct interactions with epithelial cells function to prevent widespread TR-AM proliferation in the presence of physiological levels of GM-CSF.^52^ Thus, it may be that previous observations of HIF-1*α* induction in TR-AMs are related to proliferation as opposed to inflammatory effector function.

While HIF-1α deletion in BMDMs depressed proinflammatory effector function, RNA-seq Gene Ontology analysis showed upregulation in ribosomal biogenesis, rRNA metabolic processes, and noncoding RNA processing gene pathways in *Hif1a^-/-^* BMDMs. Others have recently observed upregulation of ribosomal biogenesis genes in tumor-associated macrophages, but the mechanism underlying this phenomenon remains unclear.^53^ Increased expression of these gene pathways has been associated with increased proliferation and cell growth, and c-Myc has been identified as a critical regulator of these processes.^24^ c-Myc and HIF-1α can work in concert or opposition to augment cellular metabolism, protein synthesis, cell growth, and proliferation.^25^ While it may seem paradoxical, *Hif1a^-/-^* tumors grow faster and become more invasive in the presence of adequate oxygen, but under hypoxia, HIF-1α can displace c-Myc from DNA to induce cell cycle arrest.^54,55^ In a similar vein, BMDMs restimulated with MCSF following a period of starvation upregulate c-Myc and shift toward a proliferative phenotype reliant on both glycolysis and mitochondrial metabolism. LPS treatment downregulates c-Myc, upregulates HIF-1α, and supports a glycolysis dominant metabolism in BMDMs.^56^ This suggests that BMDMs proinflammatory effector function suppresses proliferation and shifts the cell from Myc-dependent to HIF-1α-dependent metabolism. We find the Myc regulon to be the most upregulated transcriptional network in *Hif1a^-/-^* BMDMs compared to controls. This suggests that c-Myc serves a more dominant role in the absence of HIF-1α. This likely explains the upregulation of ETC protein expression and pathways involved in ribosomal biogenesis in our *Hif1a^-/-^* BMDMs. Without HIF-1*α*, c-Myc can serve a larger role in shaping BMDM metabolism, which fundamentally changes how BMDMs differentiate and respond to immune stimuli.

In conclusion, HIF-1α is not required for TR-AM function under steady-state conditions or for inflammatory effector function, and only hypoxic conditions necessitate HIF-1α glycolytic reprogramming in TR-AMs to ensure optimal cellular fitness. HIF-1α functions much differently in BMDMs. HIF-1α is required for basal glycolytic metabolism and optimal proinflammatory effector function in BMDMs but is dispensable for LPS-induced glycolytic flux. Moreover, the loss of HIF-1α in BMDMs enhances mitochondrial function potentially in a c-Myc-dependent manner. These data describe a divergent role for HIF-1α in primary macrophage subsets and may be beneficial in developing therapies for ARDS where TR-AMs function to alleviate disease while infiltrating, nonresident macrophages exacerbate disease.

## Acknowledgments

This work was supported by funding from the Department of Defense grant HT9425-24-1-0138 (GMM), and from NIH grants R01HL151680 (RBH), F32HL167569 (ORS), R01ES015024 (GMM), and T32HL007605 (GMM).

## Author contributions

P.S.W., R.B.H, G.M.M. conceptualized and designed the study. P.S.W., R.C.A., A.Y.M., K.A.S., O.R.S., K.W.D.S., Y.F., B.H. conducted the experiments. P.S.W., A.Y.M., K.A.S., R.B.H, G.M.M. performed data analysis. R.C.A. and K.W.D.S. analyzed the sequencing data. P.S.W., R.B.H, G.M.M. wrote the initial draft, with all other authors providing comments. P.S.W., R.C.A., R.B.H, G.M.M. edited the manuscript. All authors read and approved the manuscript.

## Supplemental information

**Figure S1.**
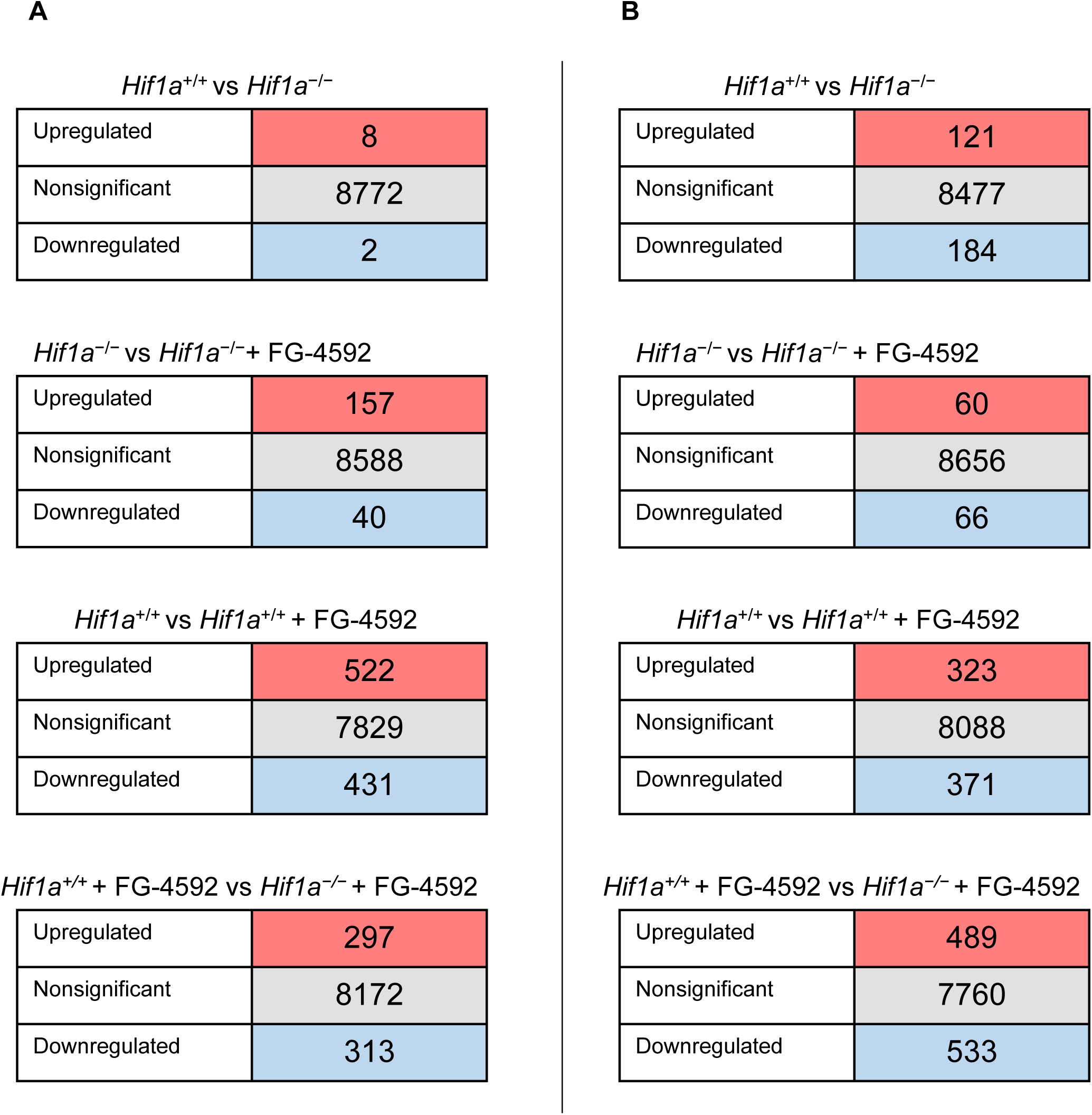
Table of DEGs related to Figure 1. **(A)** TR-AMs and **(B)** BMDMs were cultured overnight (16h) in the presence or absence of FG-4592 (25µM). Gene tables derived from RNA-seq analysis.

**Figure S2.**
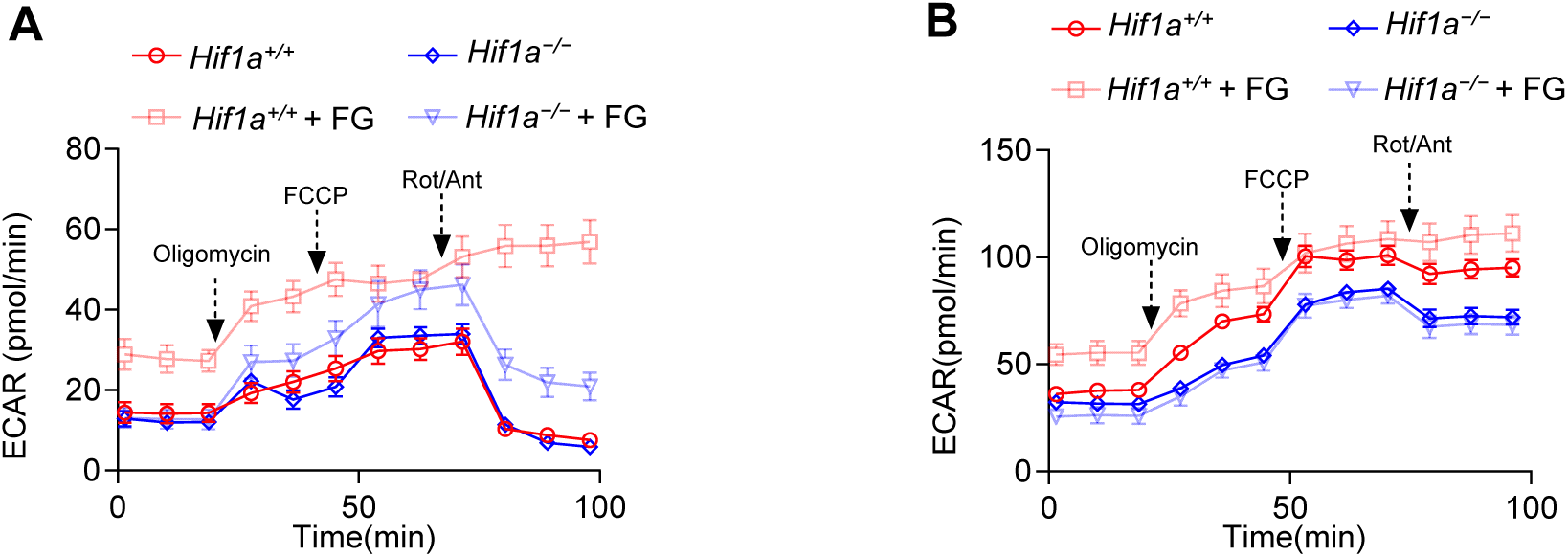
Reciprocal ECAR measurements of OCRs, related to Figure 3. Reciprocal ECAR measurements in **(A)** TR-AMs and **(B)** BMDMs during mitochondrial stress test to assess glycolytic capabilities in response to mitochondrial inhibition.

**Figure S3.**
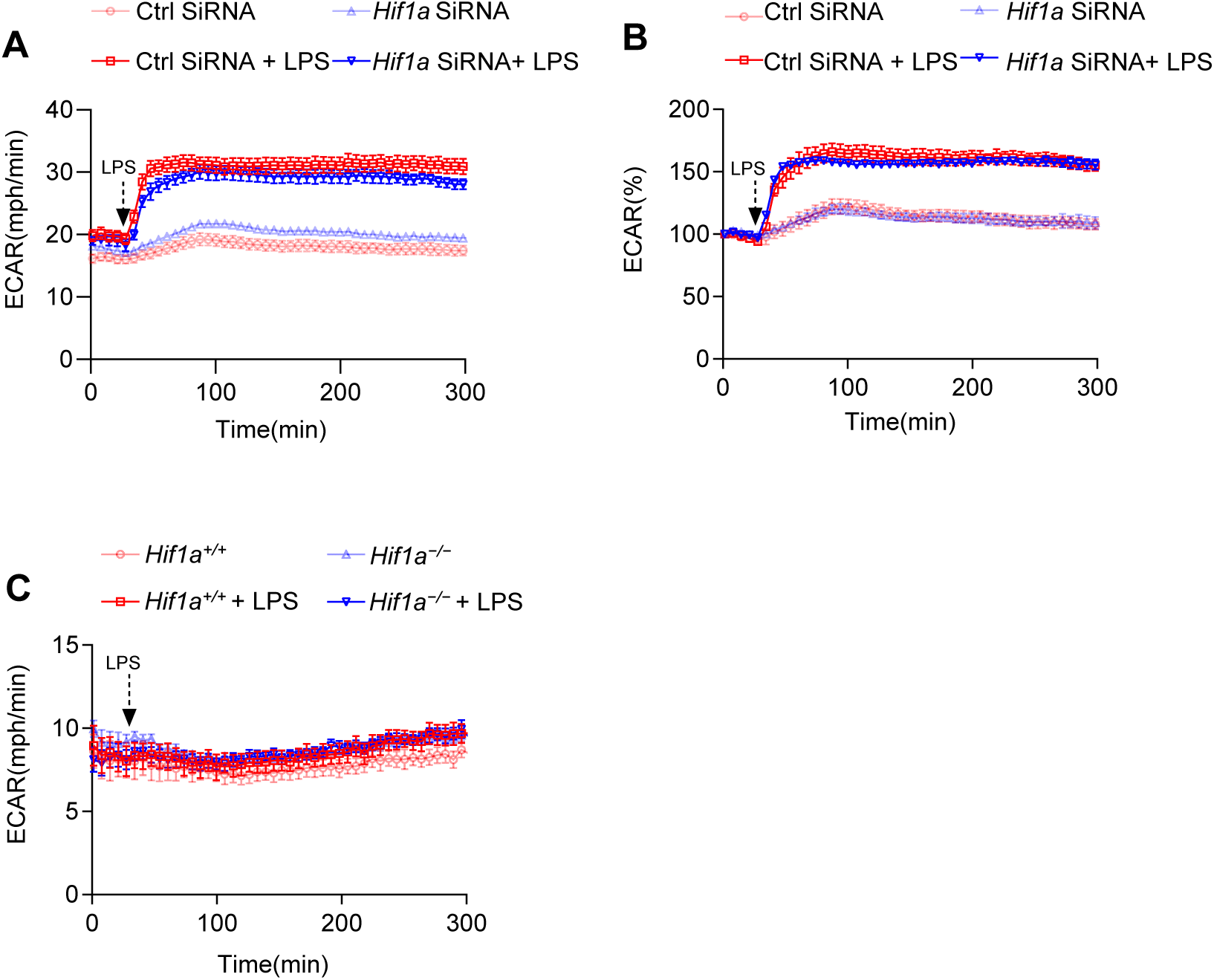
Seahorse data, related to Figure 5. **(A,B)** BMDMs were electroporated with non-targeting (Ctrl) or HIF-1α SiRNA. Cells were allowed to rest for 48 hours then subjected to seahorse analysis**. (A, B)** BMDMs and **(C)** TR-AM ECAR was measured following acute LPS injection (final concentration: 20 ng/ml). ECAR data represented both as **(A)** raw values and **(B)** % change from baseline. Data represent as at least 3 independent experiments (n=4 separate wells per group).

**Figure S4.**
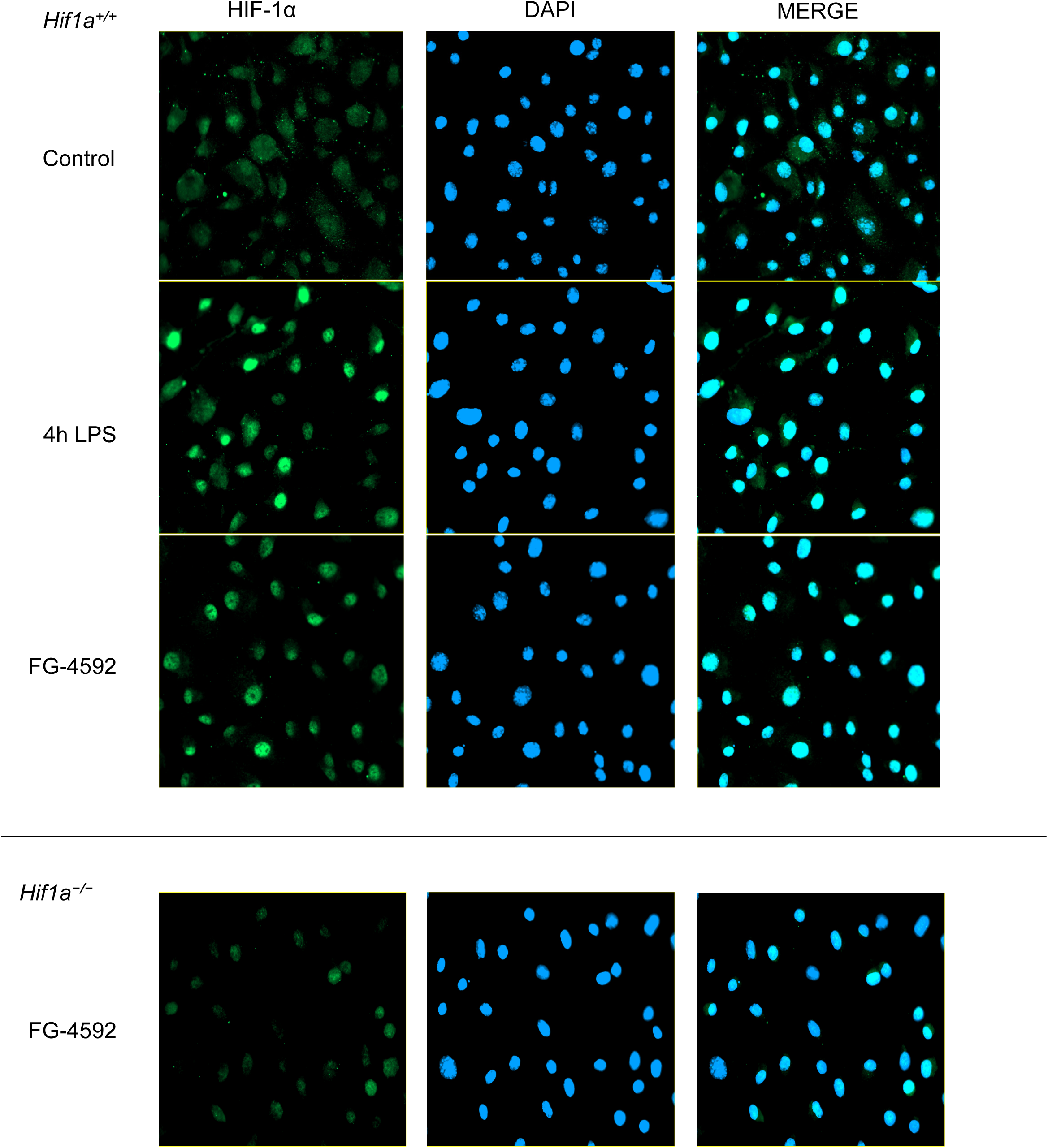
Microscopy Images, related to Figure 5. BMDMs were treated with FG-4592 (25µM) or LPS (20ng/ml) for 4 hours to assess HIF-1⍺ localization via immunofluorescence. *Hif1a^−/−^* BMDMs treated with FG-4592 served as negative controls.

**Figure S5.**
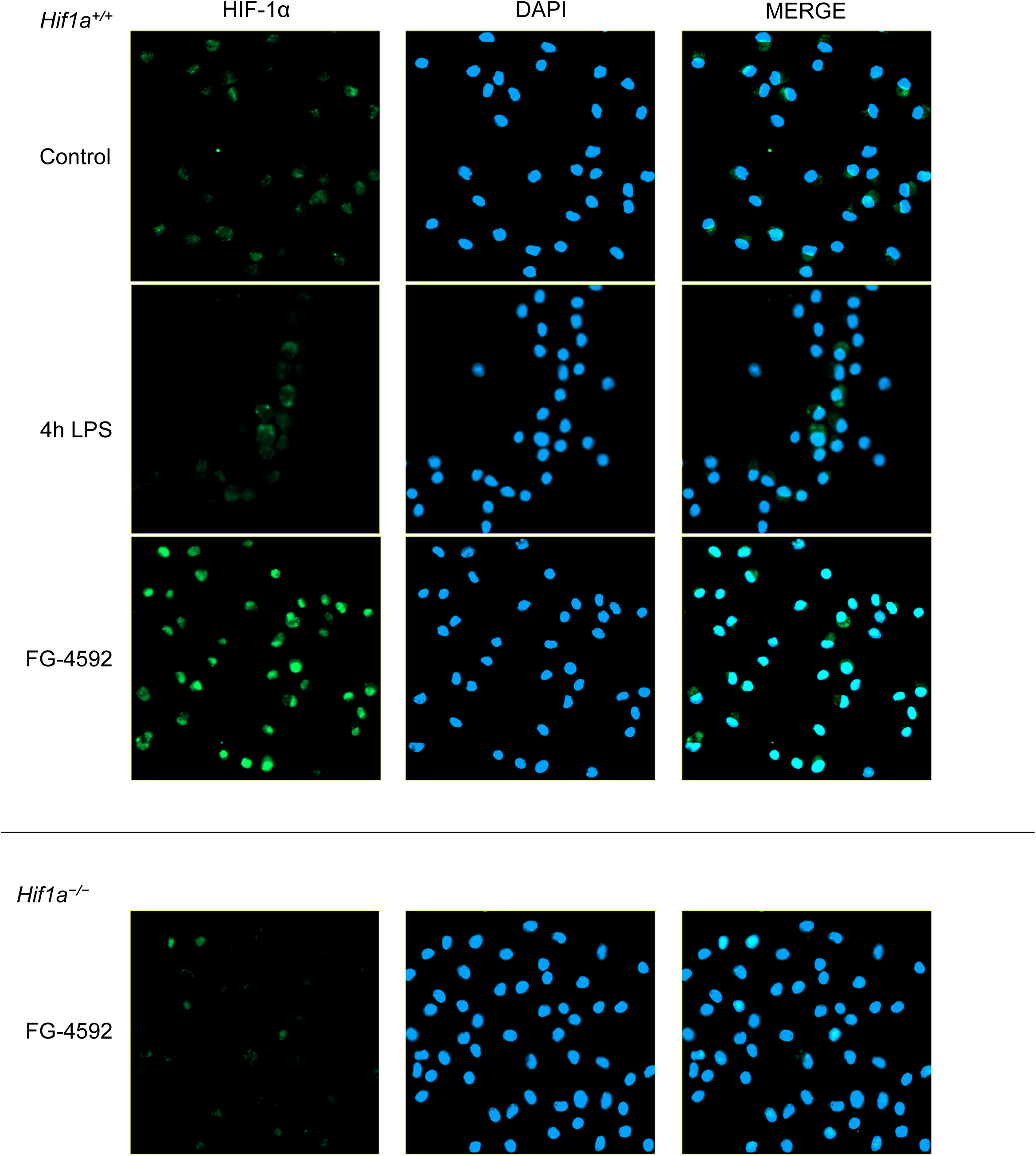
Microscopy Images, related to Figure 5. TR-AMs were treated with FG-4592 (25µM) or LPS (20ng/ml) for 4 hours to assess HIF-1⍺ localization via immunofluorescence. *Hif1a^−/−^* TR-AMs treated with FG-4592 served as negative controls.

**Figure S6.**
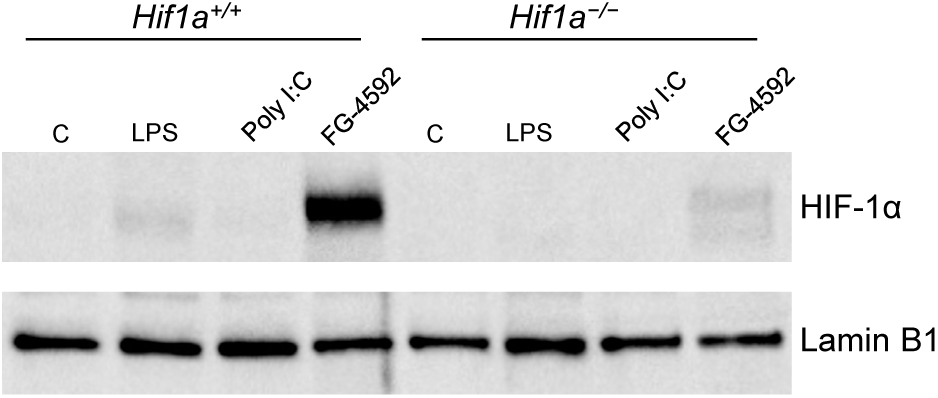
Western blot, related to Figure 5. TR-AMs were treated with FG-4592 (25µM) or LPS (20ng/ml) for 4 hours, or Poly I:C (5ug/ml) for 24 hours to assess HIF1⍺ localization expression via western blot. *Hif1a^−/−^* TR-AMs served as negative controls.

**Figure S7.**
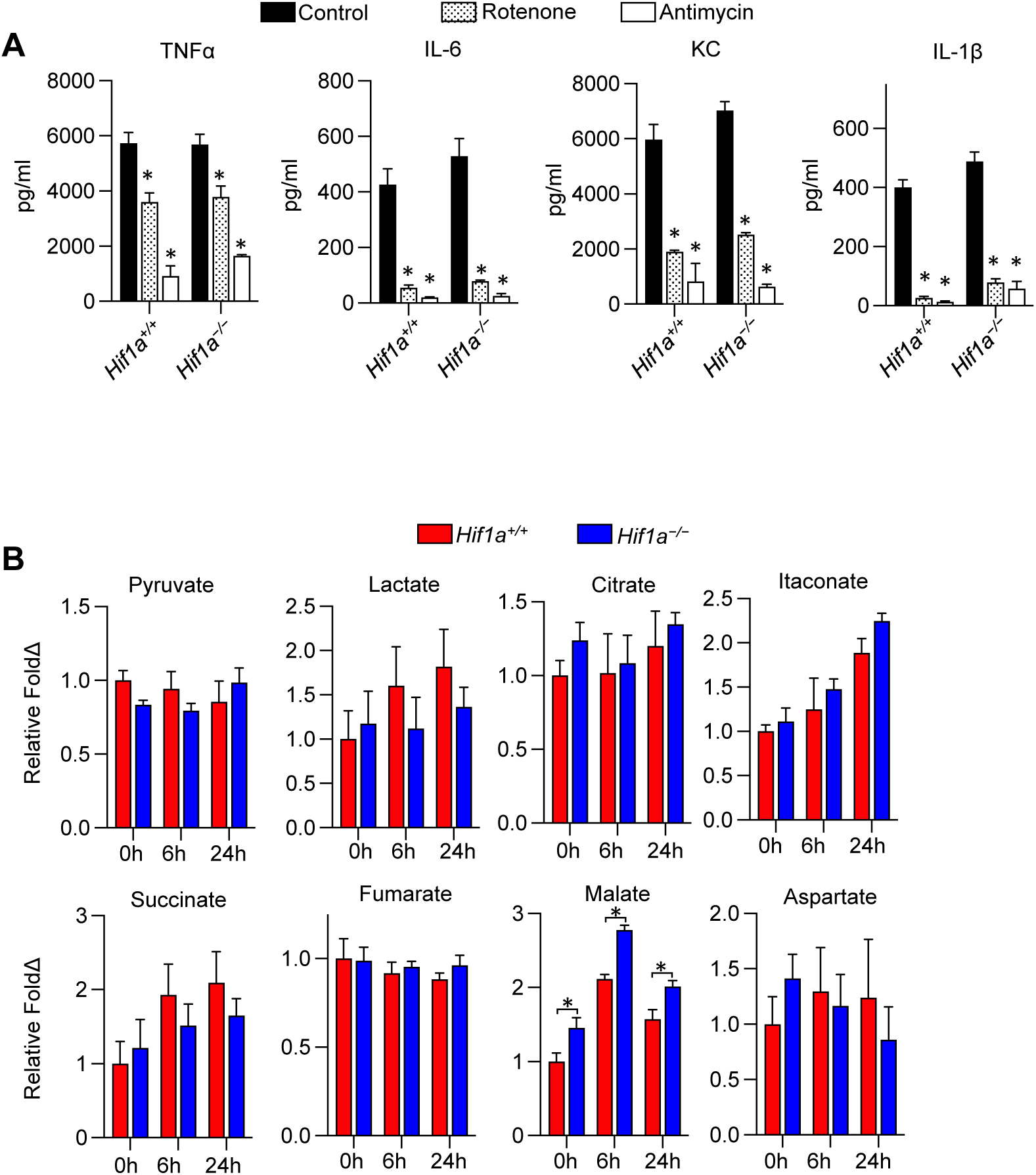
ELISA and GCMS, related to Figure 5. **(A)** TR-AMs treated with LPS (20ng/ml) in the presence or absence of Antimycin A (20nM) or Rotenone (20nM) for 6 hours. Significance determined by two-way ANOVA with Bonferroni’s post test. All error bars denote mean ± SD. Significance markings represented in relation to *Hif1a^+/+^* control group with *, p < 0.05. **(E)** GC-MS metabolite analysis of TR-AM LPS (20ng/ml) time course. Significance determined by two-way ANOVA with Bonferroni’s post test. All error bars denote mean ± SD. Significance markings represented in relation to *Hif1a^+/+^ vs Hif1a^−/−^* at a given timepoint with *, p < 0.05.

## Methods

### Primary Culture of Macrophages

All studies in animals were approved by the Institutional Animal Care and Use Committee at the University of Chicago IACUC. All animal studies were carried out according to the recommendations from an NIH-sponsored workshop ^57^ and ARRIVE guidelines^58^. To generate mice with inducible HIF1α knockout (*Hif1a^-/-^*) in cells of myeloid lineage, *Lyz2^tm1(cre/ERT2)Grtn/^*^J^ (Jackson Strain #: 031674) mice were bred with *B6.129-Hif1^atm3Rsjo/J^* (Jackson Strain #: 007561) mice. The resulting HIF1α:LysM-Cre (*Hif1a^ΔLysM^* or *Hif1a^-/-^)* and HIF1α:LysM-NoCre (*Hif1a^fl/fl^* or *Hif1a^+/+^*) progeny were treated intraperitonially with tamoxifen dissolved in peanut oil for 5 days (80mg/kg/day) to generate *Hif1a^-/-^* TR-AMs and tamoxifen-treated control TR-AMs, respectively. These 6-8 week old male mice were humanely euthanized, and their TR-AMs were isolated via standard bronchoalveolar lavage (intratracheal instillation) using PBS + 0.5 mM EDTA. Following isolation, TR-AMs were counted, plated in RPMI 1640 (ThermoFisher, Cat# 11875119) supplemented with 10% FBS (Gemini, Cat# 100-106) and 1% penicillin-streptomycin (Gemini, Cat# 400-109), and allowed to adhere to tissue culture plates for one hour prior to experimentation. BMDMs were generated by isolating bone marrow cells from the femur and tibia bones of 6-8 week old *Hif1a^ΔLysM^* (*Hif1a^-/-^*) and *Hif1a^fl/fl^* (*Hif1a^+/+^*) mice. Bone marrow cells were differentiated into BMDMs on petri dishes using 40 ng/mL recombinant M-CSF (BioLegend, Cat# 576406) and 1µg/ml 4-Hydroxytamoxifen (Millipore Sigma, Cat# SML1666) in the same media formulation as TR-AMs. On day seven, after successful differentiation and gene deletion, BMDMs were replated and allowed to adhere to tissue culture plates for two hours prior to experimentation. After adherence, cells were washed, and placed with fresh media under experimental conditions. FG-4592, a prolyl hydroxylase inhibitor, was used at a concentration of 25µM for HIF-1α stabilization. Lyophilized FG-4592 was dissolved in DMSO and diluted 1:2500 in media to achieve a final concentration of 25µM. DMSO diluted to 1:2500 in media served as untreated control groups. For inflammatory stimulation, LPS was used at a concentration of 20ng/ml.

### Bioenergetic Measurements

Glycolytic and mitochondrial respiration rates were measured using the XFe24 Extracellular Flux Analyzer (Agilent, Santa Clara, MA). BMDMs and TR-AMs were seeded at 4.0 × 10^4^/well onto Seahorse XF24 Cell Culture Microplates. Cells were equilibrated with XF Base media (Agilent, Cat# 103334-100) at 37 °C for 30 minutes in the absence of C02. Glycolytic rate was assessed using the manufacturers’ protocol for the Seahorse XF Glycolysis Stress Test followed by sequential injections with glucose (10mM), oligomycin (1.0μM), and 2-DG (100mM). Mitochondrial respiration rate was measured using the Seahorse XF Mito Stress Test according to the manufacturer’s protocol followed by sequential injections with oligomycin (1.0μM), FCCP (1.0μM for BMDMs and 4.0μM for TR-AMs), and rotenone/antimycin A (1.0μM). Assessment of real-time metabolic responses to LPS was performed using the protocol detailed in an application note provided by the Agilent ^22^. In brief, following plating, cells were equilibrated in XF base media supplemented with 10 mM glucose, 2 mM L-glutamine, 1 mM sodium pyruvate (Sigma, Cat# 11360070) and 5 mM HEPES (Sigma, Cat# 15630080), pH 7.4 and incubated at 37 °C without CO_2_ for 30 minutes prior to XF assay. Baseline metabolic rates were measured followed by direct injection of LPS (final concentration:20ng/ml). Bioenergetic rates were subsequently measured every three minutes for approximately 5 hours in total.

### Cell lysis, subcellular fractionalization and Immunoblotting

Whole cell lysates were prepared by scraping cells into lysis buffer containing 25mM Tris•HCl (pH 7.6), 150mM NaCl, 1% NP-40, 1% sodium deoxycholate, 0.1% SDS, 0.1% Benzonase, and Halt™ Protease Inhibitor Cocktail (ThermoFisher, Cat# 78430). Samples were centrifuged at 16,000 x *g* at 4 °C for 5 min to pellet cellular debris. Subcellular fractionalization and lysate preparation were carried out using the NE-PER Nuclear and Cytoplasmic Extraction Reagents (ThermoFisher, Cat# 78833). Lysate protein concentration was determined using the Pierce™ BCA Protein Assay Kit (ThermoFisher, Cat# 23225). Samples were heated to 95°C for 5 minutes and equal concentrations of samples (15μg for whole cell lysates and 5μg for nuclear fractions) were resolved on Criterion 4-20% gels (Bio-Rad, Cat# 5671093, and 5671094) and transferred to nitrocellulose (Bio-Rad, Cat# 1620167). Primary antibodies used were rabbit anti-HK2 (Cell Signaling, Cat# C64G5, 1:1000), rabbit anti-LDHA (Cell Signaling, Cat# 20125, 1:1000), rabbit anti-PHD2/Egln1 (Cell Signaling, Cat# 4835, 1:1000), rabbit anti-MCT4 (Proteintech, Cat# 22787-1-AP), rabbit anti-Lamin B1 (Proteintech, Cat# 12987-1-AP, 1:1000), rabbit anti-HIF-1α (Caymen Chemical, Cat# 10006421, 1:500), rabbit anti-c-Myc (Abcam, Cat# ab32072), and rabbit anti-*⍺*-tubulin (Proteintech, Cat# 11224-1-AP, 1:2,000). Secondary antibodies used were anti-rabbit IgG HRP-linked antibody (Cell Signaling, Cat# 7074, 1:2,500) and anti-mouse IgG HRP-linked antibody (Cell Signaling, Cat# 7076, 1:2,500). Protein expression was visualized using Immobilon ECL Ultra Western HRP Substrate (Millipore Sigma, Cat# WBULS0500) in combination with the BioRad ChemiDoc Touch Imaging system. All immunoblot data were repeated in at least three independent experiments.

### Total OxPhos Immunoblotting

Whole cell lysates were collected and quantified as described in the previous methods section. Samples were heated to 37°C for 30 minutes and equal concentrations of samples (15μg) were resolved on Criterion 12% gels (Bio-Rad, Cat# 5671044) and transferred to PVDF (Bio-Rad, Cat# 1620177). Mitochondrial complex proteins were resolved using mouse anti-OXPHOS Rodent WB Antibody cocktail (Abcam, Cat# ab110413, 1:1000).

### SiRNA knockdown

SiRNA knockdown was performed using the Amaxa Mouse Macrophage Nucleofector Kit (Lonzo, Cat# VPA-1009). 1.0 × 106 cells/reaction were resuspended in transfection solution with siRNA of interest (Dharmacon, Non-Targeting Control siRNA: D-001810-01; mouse Hif1a siRNA #1; J-040638-06). The cell solution was then subjected to electroporation (Lonza Nucleofector 2b Electroporator: Setting Y-001). Cells were plated and allowed to rest for 48 hours prior to further experimentation.

### Cytokine Analysis

Secreted TNFα, IL-6, KC, and IL-1β levels were evaluated in macrophage media using a standard sandwich ELISA (R&D Systems DuoSet ELISA Development System, Cat# DY410, DY406, DY453, and DY401). For IL-1β sample collection, 5mM ATP was added to macrophage cultures for 30 minutes following 6h LPS treatment to activate caspase 1, ensuring proIL-1β cleavage and IL-1β release. Rotenone and Antimycin A concentrations were 20nM when used in ELISA experiments.

### Sulforhodamine B (SRB) Colorimetric Assay

In vitro cytotoxicity was measured using the SRB assay ^59^. Following treatment, cells were fixed in 10% TCA and then stained with SRB dye. Cellular protein-dye complexes were solubilized in 10mM Tris base and the samples were read at OD 510 using a microplate reader. Data was normalized to the untreated, *Hif1a^+/+^* groups, which were representative of no cellular damage. ETC inhibitor concentrations were as follows: 500nM rotenone, and 500nM antimycin.

### Immunofluorescence

Macrophages were plated at 4.0 × 10^4^ cells in 50µl of media on chamber slides (ThermoFisher, Cat# 177402PK) to prevent cell dispersion toward the chamber edges. Once adhered, an additional 150µl of media containing treatment conditions (Final concentration: 20ng/ml LPS or 25µM FG-4592) was added to the chamber slides. After 4 hours, the media was carefully removed with a micropipette (no vacuum suction) and the cells were fixed in 4% paraformaldehyde for 20 minutes at room temperature (RT). Treatments were handled in this fashion because the macrophages peel off the glass slide with little agitation prior to fixation. Following fixation, cells were permealized in blocking solution (0.1% Triton X100, 3% FBS in PBS) for 60 minutes at RT. Both primary rabbit anti-HIF1*⍺* antibody (Abcam, Cat# ab179483, 1:100) and secondary Goat anti-rabbit CoraLite®488-Conjugated (Proteintech, Cat# SA00013-2, 1:400) incubations took place overnight at 4°C in a humidifying chamber. Coverslips were then mounted ProLong™ Glass Antifade Mountant with NucBlue™ Stain (ThermoFisher, Cat# P36983). Fluorescent mages were obtained using the Zeiss Axio Observer 7 Microscope. Images were prepared for publication using QuPath bioimaging software ^60^. The same brightness and contrast settings were maintained across all sample images to ensure accurate comparison.

### Gas Chromatography Mass Spectrometry (GC-MS) for metabolite measurements

Macrophages were plated at 2.5 × 10^5^ on 24-well plates for metabolite extraction. Following treatment, cells were washed with ice-cold blood bank saline (ThermoFisher, Cat# 23-293-184) and cells were scrapped into 600μl of 80% methanol with 1μg Norvaline/600μl. The metabolite samples were vortexed and centrifuged at 16,000 x *g* at 4 °C for 10 min to precipitate insoluble material. 400μl of the liquid sample was then dried under nitrogen gas for approximately 2 hours until no liquid remained. Samples then underwent derivatization, a process involving the addition of chemical modifiers to the metabolites for a more sensitive and accurate identification. First, samples were incubated in 16μl Methoxamine (MOX) Reagent (ThermoFisher, Cat# TS45950) for 1 hour at 37°C. Samples were then incubated for 1 hour at 60°C following the addition of 20ul of tert-Butyldimethylsilyl (tBDMs) (Sigma, Cat# 394882). Derivatized samples were analyzed with an 8890 gas chromatograph with an HP-5MS column (Agilent) coupled with a 5977B Mass Selective Detector mass spectrometer (Agilent). Helium was used as the carrier gas at a flow rate of 1.2ml/min. One microliter of each sample was injected in split mode (1:4 for BMDMs; 1:2 for TR-AMs) at 280°C. After injection, the GC oven was held at 100°C for 1 min and increased to 300°C at 3.5°C/min. The oven was then ramped to 320°C at 20°C/min and held for 5 minutes. The MS system was operated under electron impact ionization at 70eV and the MS source was operated at 230°C and quadrupole at 150°C. The detector was used in scanning mode, and the scanned ion range was 100-650 *m/z*. Peak ion chromatograms for metabolites of interest were extracted at their specific *m/z* with Mass Hunter Quantitative Analysis software (Agilent Technologies). Ions used for quantification of metabolite levels were as follows: Pyruvate *m/z* 174, Lactate *m/z* 233, Citrate *m/z* 591, Itaconate *m/z* 303, Succinate *m/z* 289, Fumarate *m/z* 287, Malate *m/z* 419, Aspartate *m/z* 390. For each sample, total ion counts for all metabolites were normalized internally to norvaline. All samples were then normalized to controls (HIF1*⍺*^fl/fl^; no treatment) for quantification and statistical comparison.

### RNA-Sequencing

Total RNA was extracted from cells using the GenElute™ Mammalian Total RNA Miniprep Kit (Millipore Sigma Cat#: RTN350). RNA quality was evaluated with a Bioanalyzer (Agilent), ensuring RIN values greater than 8.5. RNA was then submitted for sequencing at the University of Chicago Genomics Core Facility using the Illumina NovaSEQ6000 sequencer (paired-end). FASTQ files were generated and assessed for quality per base sequence using FastQC. RNA-seq data is accessible via GEO (GSE279117). RNA-seq reads were pseudoaligned with Kallisto v.0.44.0 at the Center for Research Informatics on the Randi high-performance computing cluster at the University of Chicago. ^61^ The Kallisto index was created with default settings using GENCODE (GRCm39), and quantification was performed in its default mode. Gene abundance calculations were conducted with the tximport R package v.1.18.0. Differential expression analysis was carried out using the edgeR R package, which involved read count filtering, normalization, dispersion estimation, and the identification of differentially expressed genes. ^62^ The library (Rtsne) was used for dimensionality reduction and visualization of variance and relationships between individual samples. Genes were considered significantly differentially expressed if they had an FDR-adjusted p-value ≤ 0.05 and a fold change (FC) greater than 2. Results were visualized with heatmaps generated from Z-score normalized expression data using the pheatmap package. Gene Ontology - Biological Process enrichment analyses plots were created using R cluster profiler package “enrichGO” function. ^63^ Genes sets used in heatmap analysis (CHEA Hif1-*⍺* target, Glycolytic and oxidative phosphorylation, c-Myc target) genes were obtained from Harmonizome, a multi-omics data integration platform.^64^ Transcription factor (TF) target gene interactions gene regulatory network enrichment analysis was done with R DoRothEA package using mouse regulons.^26^

### Statistics

The data were analyzed in Prism 10 (GraphPad Software Inc.). All data are shown as mean ± SD unless otherwise specified. ANOVA was used for statistical analyses of data sets containing more than two groups, and Bonferroni’s *post hoc* test was used to explore individual differences. Statistical significance was defined as *P* < 0.05.

